# Conjunctive vector coding and place coding in hippocampus share a common directional signal

**DOI:** 10.1101/2023.06.02.543340

**Authors:** Yue-Qing Zhou, Vyash Puliyadi, Xiaojing Chen, Joonhee Leo Lee, Lan-Yuan Zhang, James J. Knierim

## Abstract

Vector coding is becoming increasingly understood as a major mechanism by which neural systems represent an animal’s location in both a global reference frame and a local, item-based reference frame. Landmark vector cells (LVCs) in the hippocampus complement classic place cells by encoding the vector relationship (angle and distance) between the individual and specific landmarks in the environment. How these properties of hippocampal principal cells interact is not known. We simultaneously recorded the activities of place cells and LVCs via in vivo calcium imaging of the CA1 region of freely moving rats during classic, cue-card rotation studies. The firing fields of place cells rotated relative to the center of the platform to follow the cue card rotation, whereas the firing fields of simultaneously recorded LVCs rotated by the same amount as the place cells, but the axis of rotation was the nearby local landmarks, not the environmental center. We identified a novel type of place cell that exhibited conjunctive coding of the classic place field properties and LVC properties. These results demonstrate the capacity of CA1 neurons to encode both world-centered spatial information and animals’ location relative to the local landmarks, with a common directional input presumably provided by the head direction cell system.

## Introduction

Place cells in the CA1 region of the hippocampus are spatially tuned to represent an animal’s location in an environment. This tuning is achieved both by triangulating the animal’s location relative to allothetic landmarks and by integrating the animal’s speed and direction of travel to continuously update an internal representation of the animal’s location; the latter is known as path integration (1, 2). Place cells (3), grid cells (4), and head direction cells (5, 6), among other functional cell types (7, 8) are postulated to contribute to the formation of this internal, “cognitive map”, which not only supports flexible navigation but also serves as a spatiotemporal framework that binds together various components of an experience and allows the experience to be stored and retrieved as an episodic memory (9).

Objects have been used to study both the mechanisms that form the spatial framework as well as the binding of nonspatial information into the map. When placed at the periphery of an environment, objects can be used as directional landmarks to set the orientation of the internally coherent map (10). Centrally located objects are less effective in this role (10). Flexible spatial navigation and adaptive behavior also require animals to embed the landmarks and other features of the environment into the world-centered cognitive map, in order to populate the map with information about what the animal experiences in the environment. In support of this binding role, objects modulate the firing pattern of place cells within their firing fields (11). For example, “misplace cells” (3) fire in a specific location only when an unexpected object or reward appears in that place, or when an expected object or reward is missing. Furthermore, inactivation or lesion of rats’ CA1 can disrupt the memory of novel objects in a recognition task (12–14).

Animals can use individual landmarks to navigate to a specific location (15, 16). Exploration of novel landmarks modulates the firing rates of CA1 pyramidal (12, 17, 18) and LEC cells (19, 20). However, those studies do not provide strong evidence that rats generate spatial representations bound to these local landmarks. The discovery in the hippocampus of landmark vector cells (LVCs), which encode the animal’s location as a vector specifying the animal’s distance and angle to discrete landmarks (18), first confirmed the hypothesis of McNaughton et al. (21) that some place cells could exhibit such properties (18, 22). The firing fields of LVCs keep the same vector relationship to one or multiple landmarks when the locations of the landmarks are changed. Furthermore, when landmarks are moved to new locations or removed entirely from the environment, a small proportion of cells in CA1 and CA3 maintain or increase their firing at the locations where the landmarks used to be, demonstrating that these cells retain a memory for the previous locations of the landmarks (18). Other spatially selective cell types that integrate landmark information into vector representations have been reported, such as object-vector cells in the medial entorhinal cortex (MEC) (23), vector trace cells in the subiculum (24), and goal-vector cells in the hippocampus (25).

Although various types of information influence the firing patterns of place cells, the orientation of the cognitive map in the hippocampus is thought to be controlled by the head direction cell system (26–29). Head direction cells increase their firing when the rats’ head is pointing in a specific, allocentric direction, regardless of the animal’s location (30, 31). Place cells and head direction cells are tightly coupled, in that rotation of a salient, distal cue causes the corresponding rotation of place fields and head direction tuning curves relative to the center of the environment (29, 32). More importantly, under conditions in which place cells and head direction cells become decoupled from any external reference frame, they remain coupled to each other (6, 32). This result demonstrates an internal coupling between these two types of cells, rather than independent control of place cells and head direction cells by external landmarks (33). In contrast to the global, world-centered firing of place cells, LVCs in the hippocampus provide location information anchored to individual landmarks. It is unknown whether the landmark vector representations derive the orientation component of their vectors from the same source as the place cell system.

To address those questions, we trained rats to forage in an open platform with varying numbers of landmarks and utilized the miniscope calcium imaging technology in freely behaving rats. We analyzed the responses of simultaneously recorded place cells and LVCs in the hippocampus when a salient cue card was rotated around the open field. The results showed that the firing fields of place cells and LVCs rotated by similar degrees relative to the center of the environment and relative to the nearby landmark, respectively, suggesting that place cells and LVCs share a common directional input to anchor their world-centered and landmark-centered firing fields.

## Results

### The characterization of place cells in freely moving rats with calcium imaging

We performed in vivo, single photon (1P) calcium imaging in rats with head-mounted miniscopes (34) as they foraged for irregularly scattered food rewards on a square platform surrounded by a circular, black curtain (Fig. 1a). A large cue card was placed on the curtain to serve as the only salient, directional cue in the environment. Most studies using in vivo calcium imaging have been conducted in mice, with only a limited number being carried out on rats using head-mounted miniscopes (35, 36). Therefore, we optimized surgery procedures, including aspiration, lens implantation, and virus injection procedures, to successfully apply in vivo calcium imaging to rats (Fig. 1c; see Methods for details). We recorded the calcium activity from 1547 CA1 neurons from 8 rats, of which 770 were defined as active cells (∼50%), and 517 (∼33%) were defined as place cells (see Methods for definitions of “active cells” and “place cells”). These results are consistent with previous electrophysiological and IEG studies in rats that estimate that the percentage of CA1 cells that have place fields in a given environment ranges from ∼33%-50%, depending on the experimental task and environment (37–40).

**Fig. 1:**
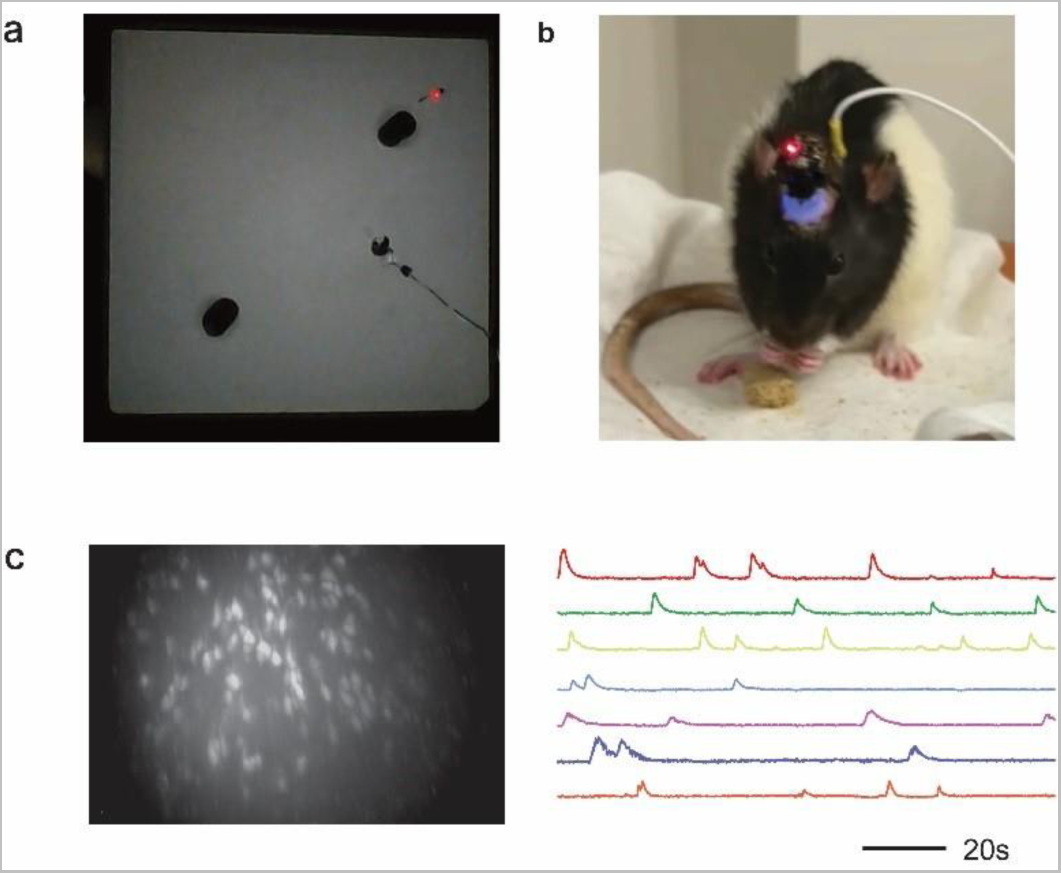
Experimental setup. **a**, Recording square platform (100 x100 cm) with 2 objects (large black spots). The recording cable and head stage are also visible. **b,** Rat with a head-mounted miniscope obtained from the UCLA open-source miniscope consortium (http://miniscope.org/index.php/Main_Page) **c**, An example of a calcium imaging video frame (left) and traces of calcium events from 7 cells (right).

### Landmark vector fields in CA1 hippocampus with calcium imaging recordings

In addition to place cells, we characterized LVCs in the calcium imaging recordings. Figure 2 shows representative examples of LVCs identified with the miniscope technique. In one experiment, as expected, the landmark vector field maintained the same direction and distance relative to the landmark when the landmark’s location was changed (Fig. 2d). In another experiment, additional landmarks were placed in an environment. In the first session, Cell 1 had a strong LV field near the landmark in quadrant 1 and a weak LV field (a single calcium transient) near the landmark in quadrant 3 (Fig. 2e, Cell 2); the latter field had the same orientation relative to the landmark in quadrant 3 as the former field relative to the landmark in quadrant 1. In the second session, in which 2 new landmarks were introduced, strong LV fields formed near both landmarks in quadrants 1 and 2, and the weak firing field was maintained near the landmark in quadrant 3; the cell did not show any detectable calcium activity near the landmark in quadrant 4 (Fig. 2e, Cell 1, right). Cell 2 formed strong LV fields in the first session near both landmarks and formed a new LV field near the landmark in quadrant 3 (Fig. 2e, Cell 2, right). The third experiment started with an open platform for the first session, in which cell 1 displayed weak, diffuse activity primarily in quadrant 3 (Fig. 2f, Cell 1, left). In the second session, two landmarks were introduced, and the cell was active at the same orientation and distance for both landmarks, along with diffuse firing at other locations in the environment (Fig. 2f, Cell 1, right). Cell 2, from the same recording session, formed a firing field near the corner of quadrant 1 (Fig. 2f, Cell 2, left); the firing field was captured by the landmark in quadrant 1 and formed another LV field near the landmark in quadrant 3 (Fig. 2f, Cell 2, right). Importantly, the examples in Fig. 2f were from the first experience that rat had with any landmarks on the platform, demonstrating that LVCs do not require experience to develop their vector fields. These results demonstrate that in vivo calcium imaging techniques can be reliably applied to identify LVCs in rats’ hippocampus.

**Fig. 2:**
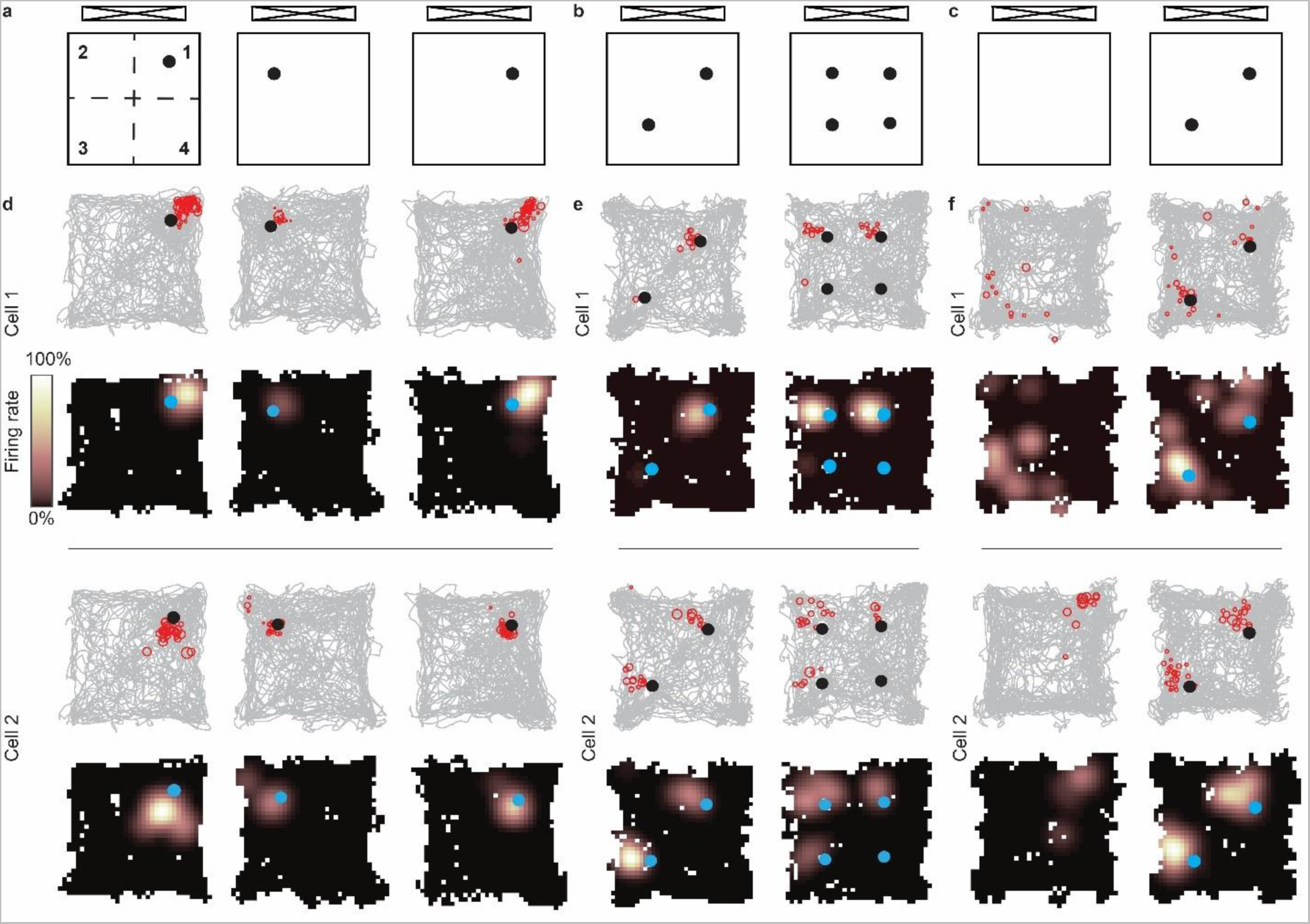
Identification of LVCs with calcium imaging recording in freely moving rats. **a-c**, Schematic of three experimental configurations. **d**, Two examples of LV fields follow the movement of the landmark. (Top) Grey: rats’ trajectory; red: locations of the rat when calcium events were detected. Black dots indicate the locations of objects. (Bottom) Occupancy-normalized calcium event rate maps; blue dots show the locations of objects. **e**, Two example cells showing additional new objects cause the addition of a corresponding LV field with the same direction and distance relative to their anchoring landmarks. **f**, Two examples of adding two landmarks to an open field causing the appearance of two LV fields. The second session was the rat’s first experience with objects on the platform. For each panel d, e, and f, the example cells were recorded simultaneously.

### Control of place cell and landmark vector cell firing fields by rotation of a salient cue card

To investigate how the distal cues affect the landmark vector representations, the location of the cue card relative to the center of the square platform was rotated between sessions (Fig. 3a), using a classic test of cue control over place cells (6, 32). In the first (standard) session, the cue was positioned north of the platform, at the same location as during prior training sessions. In the second (rotation) session, the card was shifted toward the west of the platform in between sessions. Two local landmarks were located in the centers of quadrants 1 and 3 of the platform. As expected, the spatially tuned firing fields of place cells tended to rotate ∼90° relative to the center of the platform when the cue card was rotated (Fig. 3b, Extended Data Fig. 3b). Importantly, LVCs also rotated their firing fields ∼90°, but the rotation was anchored to the nearby local landmark instead of the center of the platform (Fig. 3c; Extended Data Fig. 3c). For example, cell 2 from rat 1 showed LV responses to both landmarks in the first session, with its firing field to the northeast of each landmark. In the rotation session, the cell now fired to the northwest of the landmark in quadrant 1 and it lost its firing field in quadrant 3 (Fig. 3c, left). In contrast, cell 2 from rat 4 showed firing fields to the southeast of both landmarks in the standard session. The firing fields rotated to the northeast of both landmarks in the rotation session (Fig. 3c, right).

**Fig. 3:**
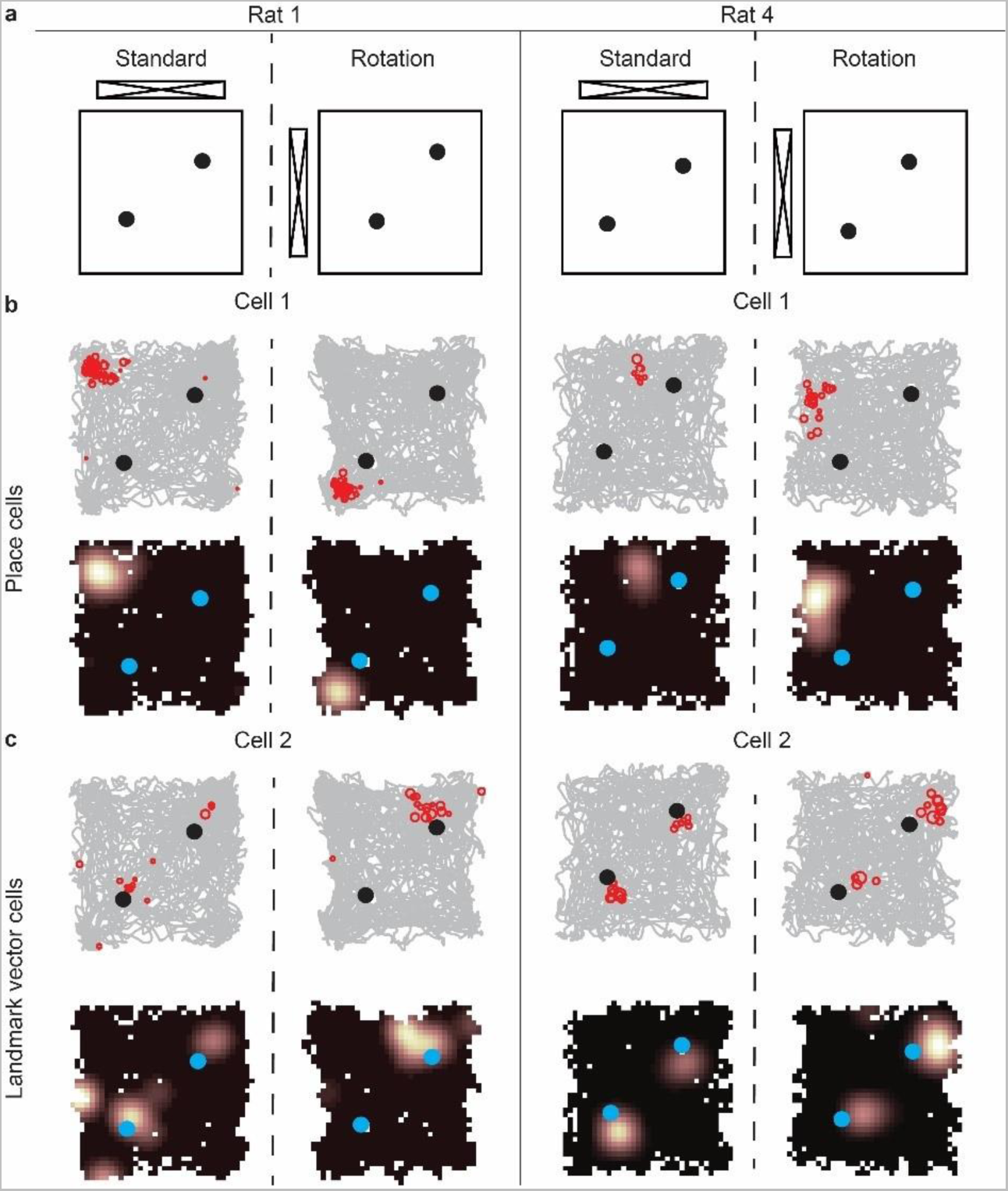
Example of place cells and LVCs response when salient cue card was rotated. **a**, Experimental configurations. **b**, Examples of place cells from two rats. The place fields rotated ∼90° relative to the center of the platform when the salient cue was rotated 90°. **c,** Examples of LVCs from two rats, recorded simultaneously with the place cells of panel **b**. Cell 2 from Rat 1 formed two LV fields near both landmarks in the standard session. The LV field anchored to the landmark in quadrant 1 rotated ∼90° in the rotation session. The field anchored to the landmark in quadrant 3 disappeared in the rotation session. Cell 2 from Rat 4 had two LV fields near both landmarks; the firing fields rotated ∼90° relative to the landmarks in the rotation session.

### Place cells and LVCs receive common directional information but are anchored to different frames of reference

We performed modified versions of the standard “rotation correlation analysis” (41–43) to quantify the degree of rotation of place fields and LV fields relative to the platform center and relative to the landmark locations. It is not always straightforward to define a field as a place field vs. a LV field, as the firing patterns can be complex and variable, especially when there are multiple landmarks in the environment and complex responses to cue rotations. We instead chose to measure the degree of rotation of all firing fields around these two axes of rotation and analyzed the population statistics of these firing fields. For rotations relative to the center of the platform, we rotated the rate map of the standard session in 1° increments and, for each increment, calculated the number of overlapping spatial bins with nonzero firing rate in both the rotated standard session and the cue manipulation session. The “platform-centered rotation degree” (PRD) was defined as the rotation angle that maximized this overlap. For rotations relative to the landmarks, we divided the rate maps into individual quadrants (Fig. 4a). Because LV fields could be anchored to different landmarks in different sessions (e.g., Extended Data Fig. 2d, Extended Data Fig. 3c,d; also (18)), we first averaged the firing rate maps of the two quadrants containing landmarks (creating “combination maps”; Fig. 4a and Extended Data Fig. 2). Because the landmarks occupied the center of the quadrants, we performed the same rotation analysis on these quadrant rate maps as for the PRD and calculated the “landmark-centered rotation degree” (LRD) as the rotation angle that maximized the overlap of nonzero firing rate bins between the standard and rotation session combination maps.

**Fig. 4:**
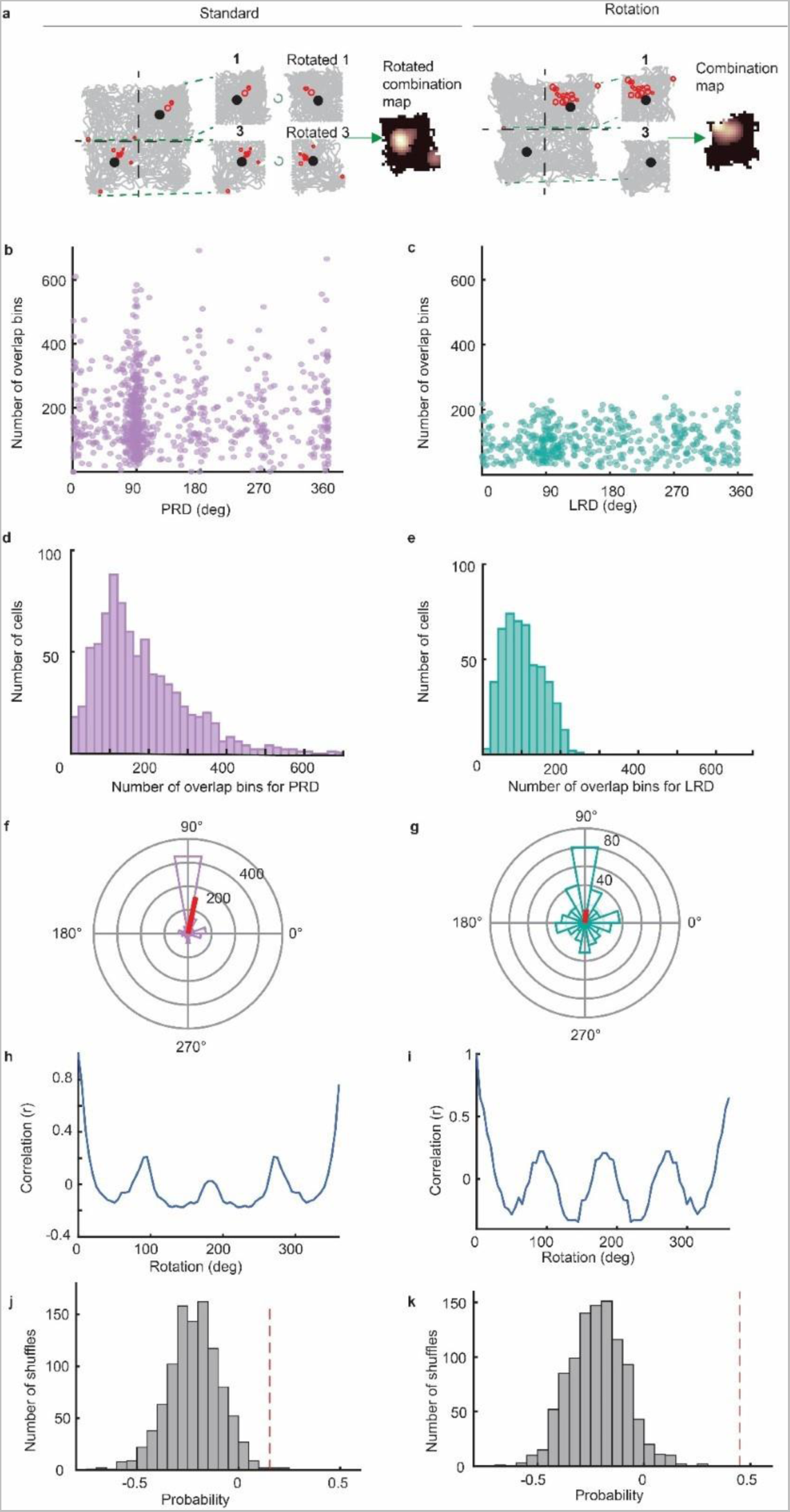
Landmark vector cell firing fields are controlled by the cue card but anchored to the nearby landmarks. **a**, Quadrants that contain landmarks (1 and 3) are incrementally rotated relative to the landmarks and merged at each increment to generate the rotated combination maps. **b**, Scatter plot shows the PRD and corresponding maximum numbers of overlap bins for all the cells. **c**, Scatter plot shows the LRD and corresponding maximum overlap bins for cells that have activity in either quadrant 1 or 3 in both sessions. **d**, The distribution of corresponding maximum overlap bins from (**b)**. **e**, The distribution of corresponding maximum numbers of overlap bins from (**c**). **f**, Polar plot of the distribution of PRD (red line denotes the direction and magnitude of the mean resultant vector: mean angle = 78.2°). **g**, Polar plot of the distribution of LRD (red line denotes the direction and magnitude of the mean resultant vector: mean angle = 83.2°). **h**, Autocorrelation plot of PRD. We rotated the polar plot of panel (**f**) in steps of 5° and computed the correlation between each rotated vector and the original vector. **i**, Autocorrelation plot of LRD, similar to that of PRD. **j**, The significance of the geometric rotation score was determined by creating a null distribution by randomly assigning a value varying from 0° to 360° to each cell and calculating the geometric rotation score for those permutations. The red dashed line indicates the actual geometric rotation score, which is greater than the null distribution (*P* < 0.05). **k**, The significance of the geometric rotation score for LRD, performed similarly to that for PRD. The red dashed line indicates the actual geometric rotation score, which is greater than the null distribution (*P* < 0.001).

On average, the degree of rotation of firing fields relative to the center of the platform equaled the degree of rotation of quadrant-based firing fields relative to the landmarks. The mean value of PRD from the 821 neurons that had detectable activity in both sessions was 78.2° ± 5.5° (Fig. 4f, r = 0.49; Rayleigh’s test, *P* < 0.001). Of these 821 neurons, 493 had activity in quadrants 1 or 3 in both sessions; the mean value of LRD from these 493 neurons was 83.2°± 20.7° (Fig. 4g, r = 0.18; Rayleigh’s test, *P* < 0.001). There was no significant difference between PRD and LRD (circular analog of Kruskal-Wallis test, *P* = 0.34). These findings suggest that place field representations and LV representations share the same directional information, each one undershooting the rotation of the cue card by ∼10° (in agreement with prior studies of head direction cells and place cells under cue-conflict situations (32, 44, 45)). The rotation of place fields around the platform center was strongly controlled by the cue card (Fig. 4f), but the distribution of LRD values was more dispersed (Fig. 4g). Part of this dispersal may be due to the smaller size of the LV firing fields compared to standard place fields, and also due to encroachment of place fields from quadrants 2 and 4 into the quadrant 1 and 3 rate maps (Extended Data Fig. 2).

The distributions of Figure 4b, c, f, and g demonstrate a 4-fold radial symmetry in PRD and LRD, suggesting that the square geometry of the environment influenced the rotation angles of the firing fields relative to the landmarks. To quantify this effect, we measured the periodicity of the PRD and LRD polar plots by computing an autocorrelation of the polar plots of Fig. 4f and Fig. 4g. The distributions of PRD and LRD were rotated in steps of 5°, and the correlation between each rotated distribution and its original was computed. If the PRD or LRD was influenced by the geometry of the platform, the values of the correlations should be greater at 90°, 180°, and 270° rotations than for 45°, 135°, 225°, and 315°. As predicted, the correlations peaked at multiples of 90° (Fig. 4h and 4i). Analogous to the standard measure used to measure the 6-fold rotational symmetry of grid cells (46), we defined the “geometric rotation score” as the minimum value of the hypothesized peak correlations (90°, 180°, 270°) minus the maximum value of the hypothesized trough correlations (45°, 135°, 225°, 315°). To simulate a null distribution, we randomly assigned an integer value from 0°–359° to each cell and calculated the geometric rotation score. The geometric rotation score of the observed data was 0.15 for PRD and was 0.45 for LRD, which exceeded the 95^th^ percentile values of the simulated null distributions, respectively (Fig. 4j, 4k; *P* < 0.05 and *P* < 0.001, respectively).

### Conjunctive representations of landmark vector fields and place fields

Since LVCs and place cells both exist in CA1 of the hippocampus and respond similarly to cue card rotations, we next investigated whether individual CA1 cells could show both classic place cell properties and LVC properties. Our results revealed a novel form of conjunctive coding of place in a subset of hippocampal place cells, in which an animal’s location is represented simultaneously in the world-centered frame (a standard place field) and also in a landmark-centered frame (a LV field) (Fig. 5; Extended Data Fig. 3d,e; see Methods). In some cases, the cell displayed distinct subfields, each of which demonstrated either place field or LV properties (Fig. 5a: Rat 1_Cell1; Fig. 5b: Rat 3_Cell1). Alternatively, cells with a single firing field observed in the standard session could be divided into two distinct firing fields when the cue card was rotated, with one field anchored to the center of the platform and the other field anchored to the landmark (Fig. 5a: Rat 1_Cell 2; Fig. 5b: Rat 3_Cell 2). Figure 5c shows a scatter plot of the distribution of PRD relative to LRD for each cell that had a valid LRD value. Collapsing the data across each axis demonstrates that the most common response was ∼90° for both measures. To address whether individual cells were more likely than chance to share this rotation amount between LRD and PRD, the difference between PRD and LRD was calculated for each cell; the distribution of this difference peaked at 0° (Fig. 5d). Of the cells that were active in quadrants that contained objects, 16% (81/493) were defined as a conjunctive place x LV cell in that the difference between the LRD and PRD for each cell was < 15°. To assess whether the population level conjunctive activity was random, we shuffled the LRD values of each cell by assigning randomly without replacement values from the distribution of LRDs to each cell. For each of the 1000 shuffles, the percentage of the cells showing conjunctive place x LV fields was computed using 4 criteria (|PRD-LRD| < 15°, |PRD-LRD| < 30°, |PRD-LRD| < 45°, and |PRD-LRD| < 65°), to show that the significance was not due to choosing a specific criterion. The percentage of the conjunctive place and LV cells in the observed data was found to be above the 95^th^ percentile of the null distribution for all criteria (Fig. 5e).

**Fig. 5:**
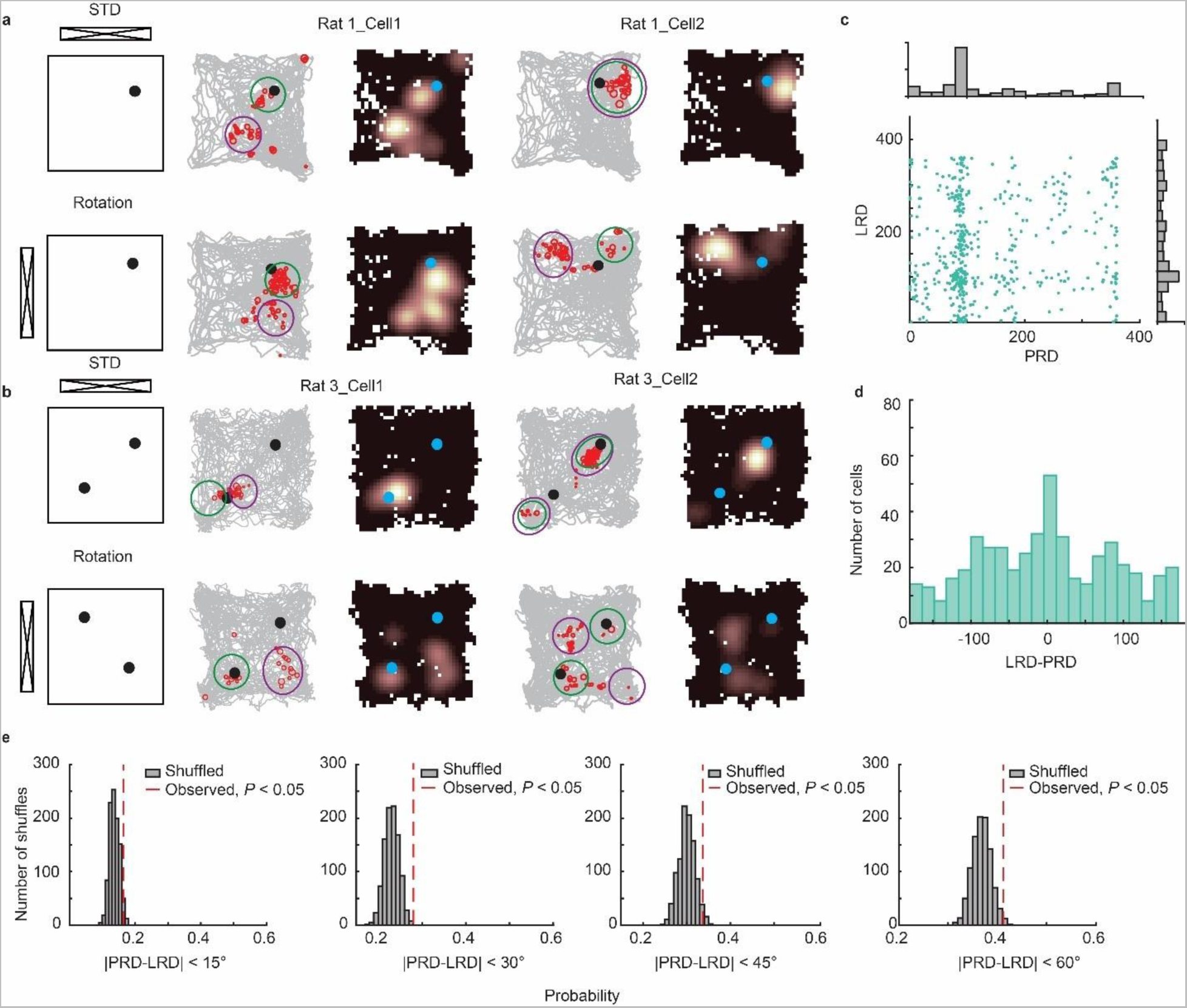
Conjunctive representations of LVCs and place fields. **a**, Example of two conjunctive representations of place and landmark vector fields when one object was on the platform. (Colored circles in all panels are meant simply to point the viewer’s attention to specific sets of calcium events corresponding to individual fields across sessions. They do not represent any quantitative measurement of field properties such as magnitude, size, etc.) Rat 1 Cell 1 had two firing fields in the standard session. In the rotation session, the green field rotated ∼90° around the landmark whereas the purple field rotated relative to the platform center. Rat 1 Cell 2 had a single place field in the standard session. In the rotation session, the firing field split into two subfields, with one (green) rotating around the landmark and the other (purple) rotating around the platform center. **b**, Example of two conjunctive representations of place and landmark vector fields when two objects were on the platform. Rat 3 Cell 1 had an elongated field near the landmark in quadrant 3 in the standard session. In the rotation session, the field split in two, as a small subfield (green) rotated around the landmark and a larger subfield (purple) rotated around the center of the platform. Rat 3 Cell 2 had 2 fields in the standard session. The two subfields broke into 4 subfields in the mismatch session, rotating around the landmark (green) and the center of the platform (purple). **c**, Scatter plot of the values of PRD and LRD for each cell. **d**, Distribution of the difference between LRD and PRD for each cell. **e**, A shuffled distribution of the difference between LRD and PRD for each cell was generated 1000 times by randomly assigning LRD values without replacement for each cell. The percentage of conjunctive representations, defined as those with |PRD-LRD| < 15° (left), |PRD-LRD| < 30° (left, middle), |PRD-LRD| < 45° (right, middle), or |PRD-LRD| < 60°, in each of those 1000 random distributions are plotted. The red dashed line indicates the observed data, which is outside the 95% confidence interval for each criterion (*P* < 0.05).

### Landmarks enhance the stability of nearby firing fields across days

Place coding in hippocampal place cells can remain stable over long periods in rats (47, 48) (but see recent work showing representational drift (48, 49)). Given that nearby landmarks offer an extra coordinate frame for animals to estimate their location, we hypothesized that nearby landmarks might increase the stability of spatial representations across days. To compare the stability between the representations near and far away from landmarks, we found that cells with fields in quadrants containing landmarks were more likely to maintain consistent activity in those quadrants across days compared to cells with signals in quadrants without landmarks. The number of active neurons in quadrant 1 or 3 on both day 1 and day n was significantly higher than the number of active neurons in quadrants without landmarks (2 and 4) on day 1 and day n (Fig. 6b; paired t-test, *P* = 0.011). Moreover, the spatial correlations of the quadrant rate maps with landmarks were higher than those without landmarks (Fig. 6c, two-sample Kolmogorov-Smirnov test, *P* = 0.014). Therefore, local landmarks enhance the stability of the nearby neural representations across days.

**Fig. 6:**
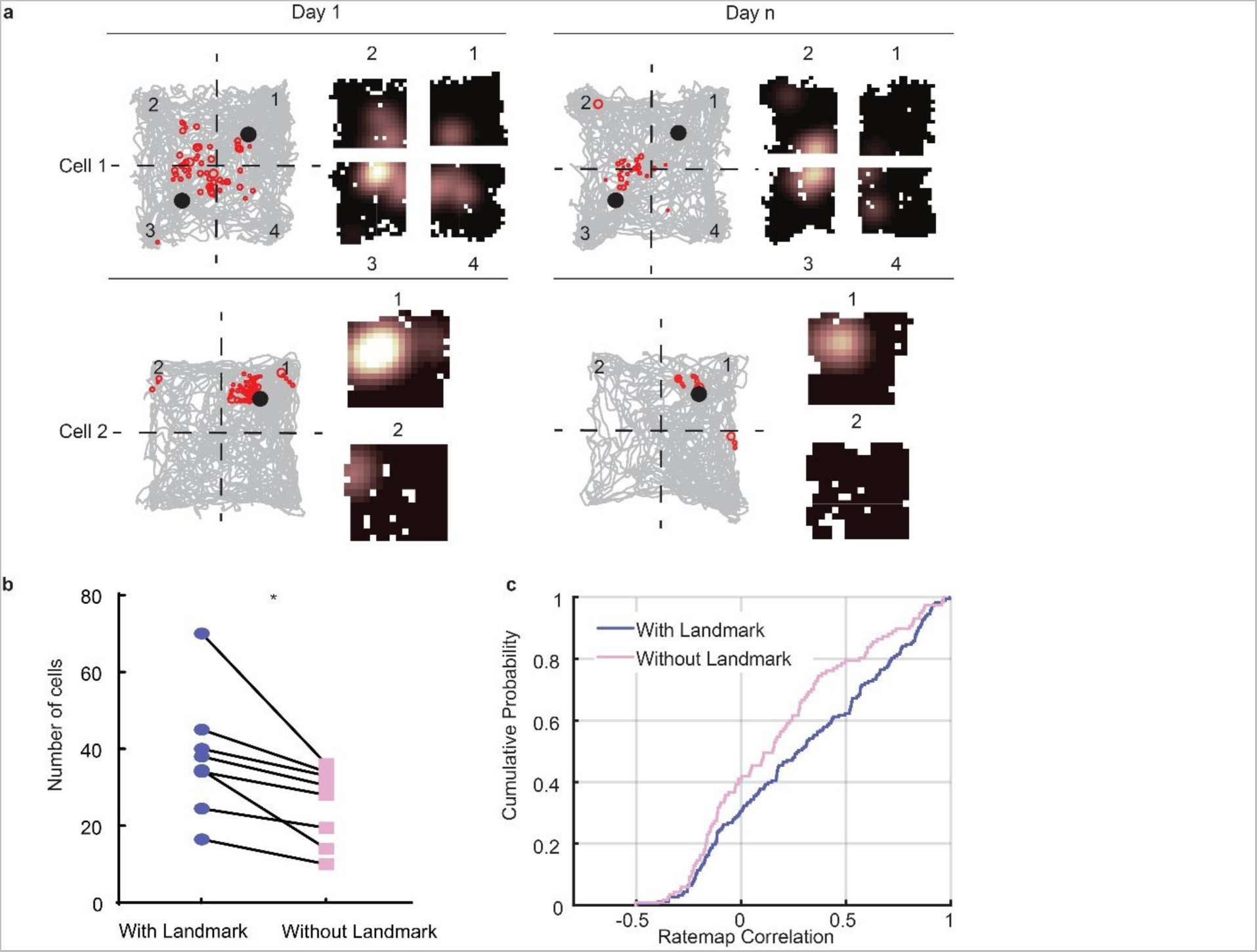
Landmarks enhance the stability of neuronal representation near the objects across days. **a**, The number of consistently active cells and the correlation coefficients between each quadrant on day 1 and its corresponding quadrant on day n were computed for all the cells. For the experimental configuration with two landmarks (top), the number of cells with consistent activity across days was calculated as the average of these values for quadrants 1 and 3 and quadrants 2 and 4. Similarly, the correlation coefficients of the quadrant rate maps were calculated as the average of these values for quadrants 1 and 3 and quadrants 2 and 4. For the experimental configuration with only one landmark (bottom), we used only quadrants 1 and 2 to calculate the numbers of consistently active cells across days and the rate map correlations across days. **b**, The number of cells exhibiting consistent responses near landmarks across multiple days is significantly greater than the number of cells showing consistent responses distant from landmarks (*P* = 0.011; paired t-test). **c**, Correlations between the quadrant rate map that contains landmarks are higher than the quadrant rate map without landmarks (*P* = 0.014, two-sample Kolmogorov-Smirnov test).

## Discussion

We recorded cells from CA1 in freely moving rats with varying numbers of landmarks in the environment to investigate how the rotation of cues affects the spatial firing properties of the landmark vector cells. In agreement with previous studies in the hippocampus (18), entorhinal cortex (23), and dentate gyrus (50), the fields of LVCs share the same vector relationship with multiple landmarks or maintain the same vector relationship relative to a landmark when the location of the landmark was changed. As expected, the orientation of the place field map was anchored to a global reference frame, as the rotation of a salient cue card in the periphery caused equivalent rotations of place fields around the center of the environment (5, 29). In contrast, the LV fields were also controlled by the cue card but were anchored to the reference frame of the individual landmarks to which they were bound. The identification of a new type of conjunctive “place x LV cell” suggests that individual neurons in CA1 have the capacity to encode both place fields and LV fields despite those two representations being anchored to two different coordinate frameworks. Finally, our results showed that the presence of landmarks enhanced the stability of nearby place fields across days.

### LVCs encode animals’ location in a landmark-centered frame

Place cells are thought to primarily encode space in an allocentric (i.e., world-centered) frame of reference (3, 28), independent of egocentric spatial information such as the orientation of animals’ body or head (51) (but see (52–54) for exceptions). The firing rates of place cells can change within their fields in the presence of landmarks (11), implicitly suggesting that place cells integrate information about landmarks in an allocentric frame of reference. However, LVCs represent landmarks in a completely different frame of reference than place cells; that is, LVCs are anchored to individual landmarks rather than to a global, world-based reference frame. The combination of representations of location in different reference frames may help the rat to evaluate the current location precisely (55), perhaps by using landmark position signals to correct for path integration errors in a manner analogous to previous proposals about the roles of boundaries in correcting such errors (33, 55–58). Landmarks may pin the internal cognitive map to a specific location in the world, and LVCs convey to downstream cells information about the rat’s location relative to the landmark in terms of direction and distance. Meanwhile, the hippocampal population activity may provide information to the downstream cells about what and/or where the landmark is in allocentric space (11, 59).

Since the LVCs and place cells encode the animals’ location in different coordinate frames, it is essential to consider whether LVCs constitute a separate class of cells from place cells or are instead a different type of response of the same class of cells as place cells. For example, distinct cell types of the MEC (e.g., grid cells and object vector cells) are considered as functionally dedicated populations of cells that are anatomically distinct from each other (23); that is, a grid cell appears to always be a grid cell in any environment under normal conditions, and it will never be an object vector cell. In contrast, the dynamic transition seen between place cells and conjunctive place x LV cells in the current study (Fig. 5; Extended Data Fig. 3) suggests that LV fields and place fields may derive from the same set of cells that sometimes act as place cells and other times as LVCs, depending on which of their inputs are active at a given moment. Regardless of whether a cell acts as a place cell or a LVC in a given environment, it appears to be controlled by a common orientation signal (8). Thus, as Deshmukh and Knierim (18) have argued previously, LVCs are better taken as descriptions of different types of responses of CA1 neurons instead of a new type of cell. In agreement with their hypothesized mnemonic roles of associating arbitrary stimuli (spatial and nonspatial) in the service of episodic memory, hippocampal cells appear to show greater flexibility in their coding properties than MEC cells, for example.

### Landmarks stabilize the spatial coding in CA1 neurons

By tracking the same cells in the same environment over multiple days, we found that the number of cells that showed consistent responses near the landmarks across days was higher than the number of cells that had activity farther away from the landmarks (Fig. 6). Furthermore, the spatial firing fields near the landmarks were more highly correlated across days than those farther away from the landmarks. These results complement a recent demonstration that population firing rates can decode the presence or absence of objects with decoding accuracy dependent on distance to the object location (59). The present results show conversely that, when the objects are present in both conditions, spatial representations near objects are more stable than representations farther from objects. The firing fields of hippocampal neurons can be stable over days in a nonchanging environment (60), but recent work has unveiled a much larger degree of “representational drift” in place cells than was previously appreciated (48, 49). Despite this representational drift, the hippocampal population still maintains the ability to represent different contexts in a sustained manner, as the parts of state space within which the representation drifts for one context remain segregated from that part of state space within which representation of the other context resides (61). Landmarks may serve as one of the contextual cues that limit the representational drift and keep the representation confined to a specific “neighborhood” of neural activity state space, preventing it from impinging on the neural neighborhood of another context. Therefore, landmarks may not only be represented in the hippocampus as individual items that occupy specific locations on the “cognitive map”, but the relative stability of these item representations may impart, via Hebbian association of coactive LVCs and place cells, added stability to the place-related representations in the hippocampus.

### The fields of LVCs bound to multiple landmarks

Although the hippocampus and MEC both contain cells that display vector coding relative to environmental landmarks, there are differences between the hippocampal landmark vector cells and the MEC object vector cells (we keep here the terminology used by Deshmukh and Knierim (18) for hippocampal cells and by Høydal et al. (23) for MEC cells). The MEC object vector cells fire in a vector relationship to all discrete landmarks in an environment (23). In contrast, hippocampal LVCs do not respond to all landmarks equally. As previously described (18), the LV representations can be bound to only a subset of the landmarks in an environment, and the LV firing fields can jump from one landmark to another in different sessions. However, contrary to previous claims (23), hippocampal LVCs do not necessarily require experience to exhibit their landmark vector properties. Deshmukh and Knierim’s (16) original report stated that, even though their strongest evidence for LVC coding came from the development of new LV fields at predicted locations with experience, it was unclear from their data whether such experience was required. The present results demonstrate that LVCs can be present from the rat’s first experience with landmarks, and thus they are similar in this regard to object vector cells of MEC. To avoid confusion between MEC and hippocampal vector coding, we advocate that all such coding in hippocampus follow the “landmark vector” nomenclature of McNaughton et al. (21), who predicted the existence of such properties in the hippocampus.

Regarding the flexibility of LVC binding to different subsets of landmarks across time, the hippocampus receives inputs from two main streams: allocentric spatial information (location, direction, and speed) from the MEC (8, 46, 62) and egocentric information about external cues and relative location from the LEC (9, 63). Like hippocampal LVCs, cells in LEC respond to multiple landmarks, but the firing is inconsistent across sessions (19). Thus, the LVC firing of place cells may be driven by the variable landmark-related activity of LEC cells as well as the stable object-vector firing of MEC cells, with hippocampal dynamics and plasticity related to both entorhinal inputs likely contributing to the differences between hippocampal inputs and outputs (64). An open question is how a downstream region knows what landmark the rat is near, if the MEC landmark vector cells fire equally at all landmarks and the LEC and hippocampal cells fire only transiently at discrete subsets of landmarks. One possibility is that the downstream areas can distinguish objects by their locations, by decoding the rat’s allocentric location from the population of place cells simultaneously firing in synchrony with the LVCs. Alternatively, it has been argued that population coding of objects in CA1 provides a greater ability to decode object identity than single-cell responses (11, 59).

### Vector navigation and vector coding in the hippocampal system

A fundamental question is whether Cartesian or polar coordinate systems are used for LVC and place field representations. Gallistel (65) has argued that, for path integration navigation, it is more efficient to represent space in a Cartesian system, and much research of place cells implicitly or explicitly assumes that the cognitive map is a Cartesian system. In contrast, O’Keefe (66) proposed a model in which spatial representations were based on a polar coordinate system anchored to the centroid of the available spatial landmarks in an environment. Under this model, place fields may be considered as a vector in a polar coordinate system, anchored to the center of the environment. Such cells are present in the postrhinal cortex (67) and perhaps LEC (68).

Recent years have witnessed a rebirth in the notion of vector navigation in rodents and vector representations in the hippocampus and related areas. Inspired by behavioral studies of goal seeking behavior controlled by vector relationships to individual landmarks (15), McNaughton et al (21) hypothesized that place cells may represent vectors to individual landmarks in an environment. Deshmukh and Knierim (18) confirmed this hypothesis, at least for a subset of CA1 place cells in a given environment that fired at a constant vector relationship with multiple landmarks in an environment. These landmark vector cells were similar to boundary vector cells previously identified in the subiculum, which fired in a vector relationship to the animal’s position relative to an extended boundary rather than a discrete landmark (7, 24, 28). Subsequently, object vector cells were discovered in the mouse medial entorhinal cortex (23). In recent years, other forms of egocentric vector cells have been discovered in rat and bat hippocampus (52, 69, 70), rat lateral entorhinal cortex (68), rat parietal cortex (71), rat postsubiculum (72), rat postrhinal cortex (67, 73), rat striatum (74), rat retrosplenial cortex (75), and human hippocampal formation (76) . These cells fire at a distance and egocentric bearing to specific items in an environment, such as landmarks, goals, boundaries, and/or the center of the environment. Place cells have also been shown to create a population code for vector navigation (25), and deep learning networks trained to perform vector navigation develop grid cell firing properties autonomously (77). Thus, the study of vector navigation and vector coding in the hippocampus and related areas is a burgeoning field that promises to reveal fundamental principles underlying the relationship between hippocampal spatial representations and navigation, as well as perhaps episodic memory (68, 78).

## EXPERIMENTAL METHODS

### Subjects

5 male and 3 female adults, Long-Evans rats (Charles River Laboratories, 4–8 months old) were successfully injected in CA1 with AAV9-GCaMP6f, implanted with a 2 mm GRIN lens, and trained to forage for food reward scattered on an open platform (Fig. 1a). Animals were individually housed in an animal facility with a reversed 12h light-dark cycle, and all recordings were performed in the dark cycle. All animal care, surgical, and housing protocols were approved by the Institutional Animal Care and Use Committee at Johns Hopkins University and complied with National Institutes of Health guidelines.

### Virus Injection

Rats were initially anesthetized with 3% isoflurane, followed by ketamine and xylazine injection; anesthesia was maintained with 0.5-1.5% isoflurane. During the surgery procedure, the depth of the anesthesia was assessed by periodic tests of responsiveness to toe or tail pinch and by monitoring breathing. Animals were placed in a stereotax, a ∼2 cm incision was made midline on the scalp, and a small craniotomy (∼0.5 mm) was made at the target coordinate (AP: -4.0; ML: 2.7). A Hamilton syringe (Hamilton, Model 1701 SN syringe, Model 1701 SN Syringe, Volume = 10 µl, Point Style: 4, Gauge: 33, Angle: 17, Needle Length: 30 mm) loaded with 3.0 µl AAV9. Syn.GCaMP6f. WPRE.SV40 virus (titer: 4 × 10^13^ GC per ml, obtained from Addgene) was lowered to a depth of 2.4 mm below the dura. Five minutes after the needle was placed at the target depth, a total of 1.5 µl virus was injected over 20 min. The syringe was left for 15 min before it was slowly withdrawn. After the virus injection, bone wax was applied to the craniotomy. Meloxicam was given subcutaneously at the end of the surgery.

### Microendoscope lens implantation

Lens implantation surgery was performed 2 weeks after the virus injection surgery. The anesthesia procedure was identical to that previously described. An incision was made on the rats’ scalp, and the periosteum was dissected from the skull. Four screws were screwed into the skull, and a 2 mm diameter craniotomy (center at AP: 4.0 mm; ML: 2.5 mm from bregma) was made. After removing the dura, the tissue was aspirated until the corpus callosum appeared, as indicated by horizontally oriented stripes of white matter axons; saline was continuously applied during the aspiration. A 32G needle was used to gently remove the horizontal stripes of the corpus callosum until the vertically oriented stripes of the alveus fully appeared in the craniotomy. A 2 mm GRIN lens (0.25 pitch, Edmund Optics) was placed at the center of the craniotomy and lowered to a depth of 2.8 mm from the skull. To increase the success rate of the surgery, we checked the quality of the calcium activity signal after the lens was temporarily secured when rats were anesthetized. The lens was cemented in place if robust signals were detected (Extended Data Fig. 1b), and neural recordings commenced two weeks later. Otherwise, a customized virus-coated lens (79) was inserted to replace the previous one and cemented in place. The lens was secured by applying cyanoacrylate glue surrounding the lens and followed with a thin layer of bone cement (PALACOS, Zimmer) on the skull. The lens was covered with Kwik-Sil (World Precision Instruments). Animals were then given dexamethasone (0.2 mg/kg, subcutaneously) and buprenorphine (0.05 mg/kg, subcutaneously) after surgery. Two to four weeks after the implantation surgery, animals were anesthetized with 2.0%-3% isoflurane, and a baseplate was attached with self-adhesive resin cement (3M) and dental cement (Coltene). A screw-secured cap covered the lens.

### Preparation of the virus coated lens

The virus-coated lenses were prepared one day before the aspiration surgery. The virus was slowly mixed with silk fibroin solution (50mg/mL, Millipore Sigma) with a ratio of 1:1 through the pipette to avoid generating bubbles. 1 µl of this mixture was applied on top of the lens surface homogeneously. After the first drop was completely dry, a second coating of 1 µl was gently applied.

### Behavior training and calcium imaging

After rats had recovered from the lens implantation surgery, they were food restricted to ∼85-90% of their free-feeding weight and trained to forage for chocolate pellets on a square open platform (100 x 100 cm) for 40 minutes per day. In most sessions, the platform contained various numbers of identical landmarks (a black cylinder, 5 cm in diameter and 20 cm high); the number varied based on experimental conditions. When present, the landmarks were placed at the centers of a subset of the platform’s 4 quadrants. Black curtains surrounded the platform, and a cue card (60 x 100 cm) was placed on the north side of the curtain to serve as the only salient orienting cue in the environment.

For the cue card rotation experiment, rats were allowed to rest for 20-25 minutes after the first session. Rats were then placed in a covered box; the experimenter rotated the box smoothly when walking between the common area of the lab and the computer room to disassociate the rats’ directional sense from that in the recording room. The covered box with the rat was taken inside the curtain by the experimenter at the entering position that keeps the same relationship with the cue card as in the previous session, with the cue card rotated.

For recordings, the miniscopes (http://miniscope.org/index.php/Main_Page) were secured to the rats’ heads, with the rat held at a position 180° opposite to the cue card location. Calcium images (752 x 480 pixels, V3 miniscope; 608 x 608 pixels, V4 miniscope; 30 Hz) were collected with the miniscope via a CMOS imaging sensor (Aptina, MT9V032C12) connected to a custom data acquisition system (DAQ). The DAQ was connected to a PC with a USB 3.0 cable and controlled by custom DAQ software. Animal behavioral data was acquired by a webcam at 30 Hz. Both imaging videos and behavioral videos were written to.avi files, and the trajectories of rats were analyzed by offline custom Python and MATLAB code to track the red LED on the CMOS of the miniscope.

### Data Analysis

Raw videos of calcium images were first concatenated, spatially downsampled by a factor of 2 (376 x 240 pixels, V3 miniscope; 304 x 304 pixels, V4 miniscope (34); http://miniscope.org/index.php/Main_Page), temporally downsampled by a factor of 4 (7.5 Hz) by custom MATLAB scripts, and motion-corrected in Matlab with NoRMCorre (80). Signal extractions, denoising, deconvolving and demixing were performed by constrained non-negative matrix factorization for microendoscope (CNMF-E) (81). To prevent the spurious detection of overlapping neurons as a single neuron, we employed a number of procedures. For each rat we chose 2-3 average cells from a single recording session and measured their diameters through image J (82). The mean diameter for each rat was fed as a parameter in CNMF-E. CNMF-E incorporates sparsity constraints that each individual neuron should be represented by only a few spatial components (83, 84). Temporal smoothing is also incorporated with CNMF-E, which assumes that the activity of a neuron changes smoothly over time to reduce the noise and help isolate different neurons. Moreover, CNMF-E uses an iterative optimization procedure to refine the spatial and temporal components over multiple iterations until convergence is achieved. Images were visually curated and any oddly shaped neurons that might reflect overlapping cells were either manually segregated into separate cells or discarded from analysis. In particular, examples of conjunctive place x landmarks cells were scrutinized to ensure that the calcium events when the rat was in the place field component came from the same cell as the events when the rat was in the LV field component. Finally, CNMF-E applied a post-processing step to remove false positives and merge overlapping neuron components. Those measures help CNMF-E accurately extract signals from individual neurons in the calcium imaging dataset.

### Tracking same cells across sessions and across days

To identify the same cells from images that are recorded from multiple recording sessions, we applied a probabilistic method to automatically register individual cells across multiple imaging sessions and determine the confidence of each registered cell (85). Initially, we applied CNMF-E analysis to produce spatial footprints for all cells captured during the first recording session. Subsequently, the same procedures were repeated for the cells imaged during other sessions within the same day or across the days. To align the footprints from the first session as a reference map and correct for translation and rotation differences across the different sessions, we computed the probability (*P*_same_) using their spatial correlation and centroid distance of a given group of cells, one from each imaging session. A group of cells were considered to have the same identity if their *P*_same_ > 0.5 (85).

### Place cell identification

The trajectories of rats were smoothed with a five-frame boxcar filter (150 ms) and filtered by velocity (3 cm/s). The square platform was binned into 2.5 x 2.5 cm spatial bins. The spatial rate map for each neuron was calculated by dividing the total number of calcium events by the animals’ total occupancy in each spatial bin. Bins with total occupancy over 0.3 sec were included in the calculation. The map was then smoothed with a 5-cm Gaussian kernel. To quantify the information content of a given neuron, the spatial information score for each neuron was calculated with the following formula (86),

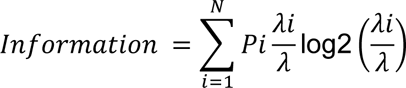

where Pi is the probability of the rat occupying i-th bin for the given neuron, λi is the neuron’s activity rate in the i-th bin (each calcium event is considered as an activity event), and λ is the overall mean rate of the neuron across the entire sessions. The timing of the calcium event was shuffled by shifting a random amount of time (minimum 50 sec) for 1000 times and the spatial information was recalculated for each shuffle. A distribution of 1000 times of shuffled spatial information was generated. A given neuron was classified as an active cell if it had a mean activity rate over 0.01 Hz; and the active cell was classified as a place cell if the observed spatial information was higher than the 95^th^ percentile of the spatial information distribution generated by 1000 shuffles (*p* < 0.05). Calcium imaging experiments in mice showed lower percentages of place cells (20%-40%) (48, 87) than those with electrophysiological studies (∼47%-70%) (48, 88, 89), whereas a previous study using miniscopes in rats reported a higher proportion (77.3%) of place cells (36). Unlike the present study, other studies with head-mounted miniscopes used the number of active cells – defined as cells that had a minimum number or rate of calcium transient activity – in the denominator to calculate place cell proportions (36, 48); this minimum number excluded cells that displayed only a single transient (or a very small number of transients), thereby underestimating the number of cells in the imaging plane. In contrast, we included all cells that fired >= 1 transient in the denominator, which, while still underestimating the denominator by excluding cells that fired no transients and were therefore invisible, produced a smaller proportion of cells with place fields that matched more closely the data from single-unit electrophysiology studies.

### Computation of LRD and PRD

To compute the landmark-based rotation degree (LRD) for each cell, we extracted the rat’s trajectory and calcium signals in quadrants 1 and 3 and created firing rate maps. We rotated the rate maps for the rotation sessions in quadrants 1 and 3 in steps of 1°, with the landmark at the center of the quadrant as the axis of rotation. The rotated combination map was generated by averaging the firing maps generated based on rotated matrices (Rotated 1 and Rotated 3; Fig. 4a, left). The number of bins in which non-zero firing rates overlapped was computed between the rotated combination map in each rotation step and the combination map in the rotation session. The LRD was defined as the peak of the smoothed number of overlap bins using the Savitzky-Golay filter (Extended Data Fig. 2f and 2i).

To calculate the platform-based rotation degree (PRD), the trajectory and calcium signals in the standard session were rotated with respect to the center of the platform in 1° increments. Subsequently, the rotated rate maps were generated based on the rotated matrices, and the numbers of overlap bins were computed between the rotated rate map in the standard session and the rate map in the rotation session. Finally, we applied the Savitzky-Golay filter to the number of overlap bins for each rotation step to identify the peak, which we defined as LRD (Extended Data Fig. 2b, 2e, and 2h).

### Geometric rotation score

Autocorrelation of the LRD distribution (5° steps) was used to measure the 4-fold, rotational symmetry of this distribution. Analogous to the standard measure of gridness for grid cells (46), the geometric rotation score was computed by taking the minimum correlation values through rotations of 90°, 180° and 270°, from which was subtracted the maximum correlation values through rotations of 45°, 135°, 225° and 315°:

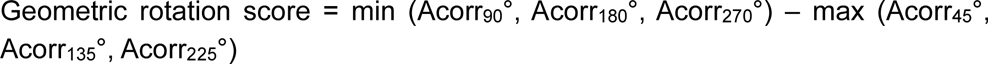

### Statistics

All statistical analyses were performed in MATLAB (Mathworks) and the results are considered statistically significant when p values are less than 0.05. Functions from the Matlab circular statistics toolbox were used to determine circular statistics (90).The circular analog of the Kruskal-Wallis test (circ_cmtest) was used to determine whether there were significant differences between the medians of PRD and LRD. The statistical procedure involving shuffles was repeated 1000 times.

### Histological Procedures

After the conclusion of the recording, rats were deeply anesthetized with Euthasol and transcardially perfused with 4% PFA prepared in PBS. The lens was carefully removed, and the extracted brain was kept in 4% PFA overnight. The extracted brain was transferred to 30% sucrose in PBS the next day. After more than 36 hours, the brain was sectioned coronally into 40-um-thick sections. Brain sections were mounted on glass slides and mounted with the fluorescence mounting medium with DAPI (Vector laboratories). Images were acquired by Zeiss LSM 780 confocal microscope. Damage to the alveus was found in a number of rats, but it was unclear whether this damage resulted from insertion of the tip of the lens into the alveus or whether it was due to healthy tissue attached to the GRIN lens being damaged during lend removal. We noticed that about half of the rats had large numbers of highly spatially tuned place cells, while the other half had many poorly selective place cells. The latter group typically showed more alveus damage than the former, and we attribute the lack of standard place cell selectivity in these rats to the hippocampal damage from the lens. Thus, we discarded all of the data from rats (n = 4) that did not show good place cells during the recordings, leaving the final number of 8 rats analyzed in the study.

## Acknowledgements

We thank Daniel Aharoni, Garrett Blair, Tad Blair, Cheng Wang, and the UCLA miniscope project for assistance with applying miniscope technology in rats. We thank Maggie Jiang for assistance with animal training. This study was funded by grants U01 NS111695 and R01 NS039456 from the U.S. Public Health Service.

## Extended data

**Extended Data Fig. 1.**
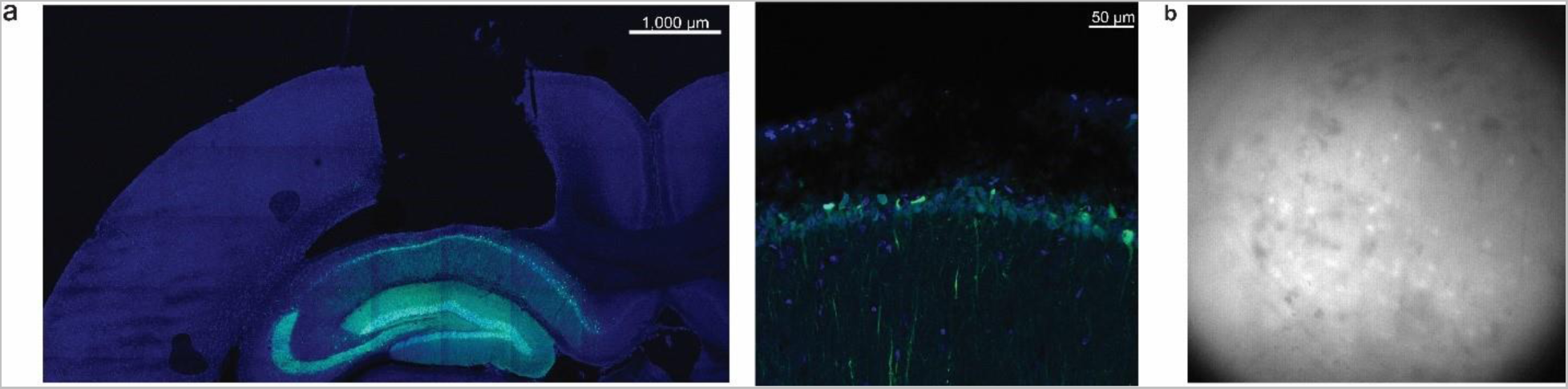
Histology of brain slice and acute signal check during the lens implantation surgery. **a**, Histology of GRIN lens implantation in HPC. Green: GCaMP6, Blue: DAPI. **b**, Example of calcium imaging raw video frame during acute signal check during the lens implantation surgery when animal was anesthetized.

**Extended Data Fig. 2.**
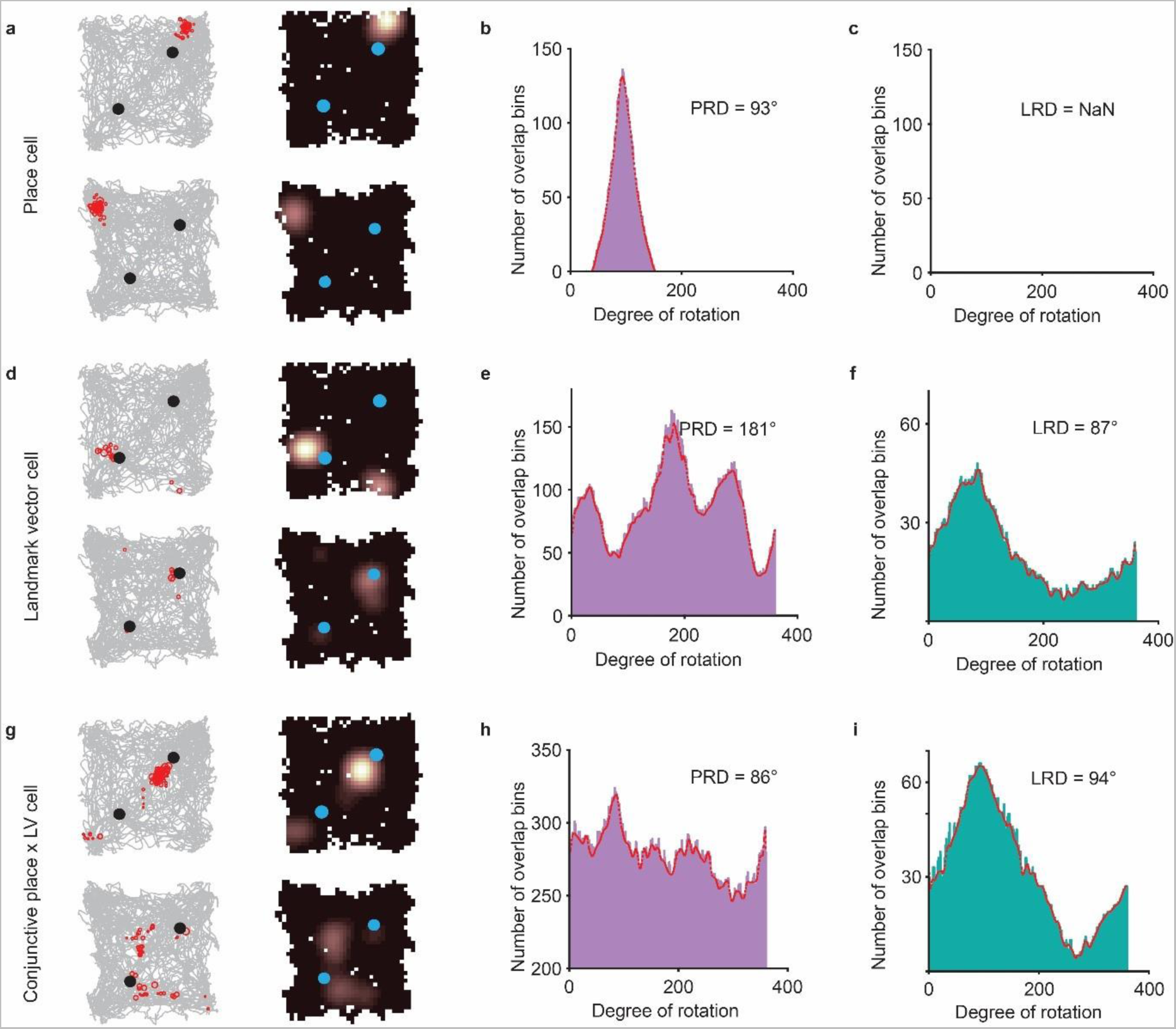
Calculation of PRD and LRD. **a**, Example of place cells in the standard trial (top) and rotation trial (bottom). **b**, The number of overlap bins between the rate maps for the standard and cue-rotation sessions peaked when the whole map in the standard session was rotated 93°; thus the LRD for the example place cell is 93°. The smoothed number of overlap bins was indicated by the red line, which was obtained through the application of the Savitzky-Golay filter. **c,** The fields of the example place cell are not in the quadrant containing landmarks during the second trial, and thus there is no LRD defined for the example place cell. **cd**, Example of landmark vector cell and the corresponding number of overlap bins when computing PRD (**e**) and LRD (**f**). The LVC rotated its firing field ∼90° (LRD = 87°) relative to the landmark, as it jumped landmarks from quadrant 1 to quadrant 3. **g**, Example of conjunctive place x LV cell. The cell rotated its firing field ∼90° in the world-centered frame (**h**, PRD = 86°) and in landmark-centered frame (**i**, LRD = 94°) when the cue card was rotated.

**Extended Data Fig. 3:**
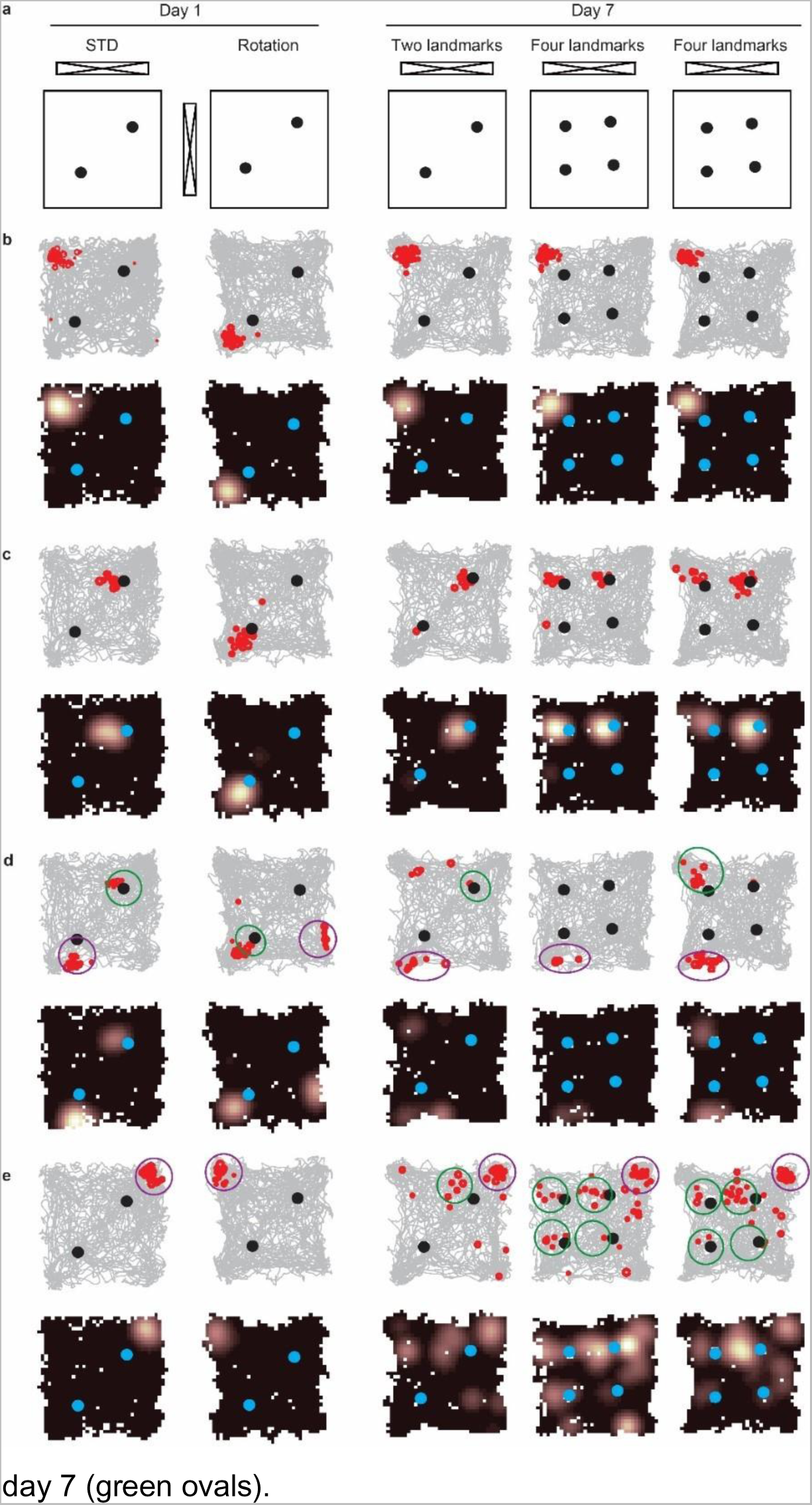
The dynamic transition of place cells and landmark vector cells for identified cells recorded across days. **a**, Experimental paradigm on day 1 and day 7. Day 1 was a cue-card rotation manipulation and day 7 introduced extra landmarks. The rat was kept in its home cage between Days 1 and 7. **b**, Example cell with a place field on day 1 that was maintained in the same location on day 7. **c**, Example LV field that rotated ∼90° relative to the landmark and jumped to another landmark when the cue card was rotated on day 1. On day 7, the landmark vector field maintained the same distance and orientation relative to the landmark and formed new LV fields when new landmarks were introduced. **d**, Conjunctive place x landmark vector cell on day 1. The LV field rotated ∼90° relative to the landmark and jumped to another landmark (green ovals) and the place field rotated ∼90° relative to the center of the platform (purple ovals). On day 7, LV fields almost disappeared in the first two trials and then strongly fired bound to the newly introduced landmark at the third trial (green ovals). The place field maintained the same position across days. **e**, Example of a cell with a place field on day 1 (purple ovals) that added landmark vector firing fields on

